# Molecular characterization of *capulet2* reveals the importance of *ANAPHASE PROMOTING COMPLEX 6* maternal expression in endosperm development

**DOI:** 10.1101/2025.09.09.675171

**Authors:** Yuri S. van Ekelenburg, Ida V. Myking, Cathal Meehan, Morten P. Hauger, Shinichiro Komaki, Keiko Sugimoto, José Gutierrez-Marcos, Paul E. Grini

## Abstract

Flowering plants are characterized by a double fertilization event, and the fertilized female gametes develop into the endosperm and embryo. Genomic imprinting promotes parental allele-specific gene expression in the endosperm by epigenetic modifications such as DNA methylation. Similarly, gametophyte maternal effects influence gene function in the female gametophyte that affects development of the endosperm and embryo post-fertilization. While most imprinted genes do not display a seed phenotype upon mutation, gametophyte maternal effect mutants are characterized by distorted seed development upon maternal transmission of the mutant allele. Here, we have investigated the gametophyte maternal effect mutant *capulet2* (*cap2*). We have established *CAP2* to be encoded by *ANAPHASE PROMOTING COMPLEX 6* (*APC6*), a subunit of the anaphase promoting complex/cyclosome (APC/C). Investigation of further *cap2/apc6* alleles revealed female gametophyte maternal effects both in mutant segregation and seed phenotype, and both *cap2* and *apc6* phenotypes were rescued by an *APC6* transgene. Furthermore, we demonstrate that *APC6* is a maternally expressed imprinted gene, in line with the observed female gametophyte maternal effect phenotype. To this end, we observed irregular nuclear division of the endosperm coenocyte in *cap2/apc6* mutants, suggesting a role for the APC/C in early endosperm development. We further demonstrate that similar endosperm defects are also produced by mutation of *APC1*, another subunit of the APC/C. Similar to *cap2/apc6, APC1* is imprinted and only expressed from the maternal allele, suggesting a maternal bias in the control of APC/C in the developing endosperm.

## Introduction

Seed development in angiosperms is initiated by a double fertilization event forming the diploid embryo and triploid endosperm (Nowack et al., 2010). The embryo develops into the next generation and receives nutrients from the mother plant through the endosperm. Due to the triploid nature of the endosperm, a balanced parental gene expression is important (Barton et al., 1984; Birchler, 1993). While most genes are expressed equally from both parental alleles (biparentally expressed genes; BEGs), some genes have been found to be regulated in a parent-of-origin dependent manner (Köhler and Grossniklaus, 2005; Nowack et al., 2010). Parent-of-origin specific gene expression, also called genomic imprinting, has been described in mammals, filamentous fungi and flowering plants (Martienssen and Colot, 2001; Feil and Berger, 2007). In genomic imprinting, an epigenetic mark is established in the gametes, resulting in silencing of the allele post-fertilization (Montgomery and Berger, 2021).

Currently, two major epigenetic regulatory mechanisms are known to be involved in the establishment of imprinting, DNA methylation and histone modification. Maternally expressed genes (MEGs) are mainly hypothesized to be regulated by DNA methylation and removal of methylated cytosines by DNA glycosylases. Several DNA methyltransferases have been identified in *Arabidopsis thaliana* (*A. thaliana*), including DNA METHYLTRANSFERASE 1 (MET1) (Finnegan et al., 1993), CHROMOMETHYLASE 3 (CMT3) (Lindroth et al., 2001) and DOMAINS REARRANGED METHYLASE 1 & 2 (DRM1/DRM2) (Cao and Jacobsen, 2002). It has been shown that DNA methylation marks established by MET1 (Xiao et al., 2003; Kinoshita et al., 2004; Gehring et al., 2006; Shirzadi et al., 2011), DRM1/DRM2 (Matzke and Mosher, 2014; Hornslien et al., 2019) and CMT3, guided by histone H3 lysine nine di-methylation (H3K9me2) (Stroud et al., 2014; Moreno-Romero et al., 2019), can regulate the imprinting state of specific genes. In addition, RNA directed de novo DNA methylation (RdDM) guided by small RNAs has been demonstrated (Vu et al., 2013; Hornslien et al., 2019; Satyaki and Gehring, 2019; Batista and Köhler, 2020).

Paternally expressed genes (PEGs) are mainly considered to be regulated by silencing of the maternal allele in the central cell by the FERTILIZATION INDEPENDENT SEED 2-Polycomb Repressive Complex 2 (FIS-PRC2) complex (Gehring, 2013). Upon recruitment of FIS-PRC2 to DNA hypomethylated loci on the maternal allele, the histone methyltransferase subunit MEDEA (MEA) trimethylates histone H3 lysine 27 (H3K27me3), a mark for gene repression (Rodrigues and Zilberman, 2015; Moreno-Romero et al., 2016). Several genes have shown to exhibit maternal silencing by the FIS-PRC2 complex, including *PHERES1* (*PHE1*), and display interaction between DNA and histone methylation (Köhler et al., 2005; Makarevich et al., 2008; Hornslien et al., 2019; Batista and Köhler, 2020). Similar to imprinted genes, gametophyte parental effect genes affect post fertilization seed development (Johnston et al., 1992; Colombo et al., 1997). Although the outcome of parental effect genes and imprinted genes is relatively similar, they are distinct genetic phenomena and have different molecular origins (Wolf and Wade, 2009). The epigenetic marks for imprinted genes are established prior to fertilization and affect gene expression post-fertilization (Montgomery and Berger, 2021), whereas parental effects are caused by pre-fertilization products (i.e. mRNA and proteins) that become active and execute their effect post-fertilization, irrespective of gene expression (Johnston et al., 1992; Colombo et al., 1997).

Gametophyte maternal effect mutants have been characterized by normal gametophyte development, followed by disrupted embryo and/or endosperm development when the mutant allele is transmitted through the female parent (Yadegari and Drews, 2004). Gametophyte maternal effect mutants have been identified in both *Arabidopsis* and maize (Evans and Kermicle, 2001; Grini et al., 2002; Olsen, 2004; Pagnussat et al., 2005; Gutiérrez-Marcos et al., 2006; Phillips and Evans, 2011; Chettoor et al., 2016), including the *capulet* (*cap*) mutants (*cap1* and *cap2*), which show developmental arrest in the early developing embryo and endosperm (Grini et al., 2002). The *cap2* allele is only lethal when transmitted maternally, indicative of a gametophytic maternal effect mutant. Gametophyte maternal effects could be explained by maternally deposited RNA or protein or by genomic imprinting. It remains technologically challenging, however, to identify the origin of transcripts in the endosperm directly after fertilization, i.e. whether transcripts come from the female central cell as maternal carry-over or if they have been transcribed de novo (Evans and Kermicle, 2001).

In this study, we have identified the molecular nature of the gametophytic maternal effect mutant *cap2*. Using whole genome sequencing and single nucleotide polymorphism (SNP) analysis, we identified the causative *cap2* SNP to be located in an intron splice donor site of the *ANAPHASE PROMOTING COMPLEX 6* (*APC6*), encoding a subunit of the ANAPHASE PROMOTING COMPLEX/CYCLOSOME (APC/C). Two independent T-DNA insertion lines, *apc6-2* and *apc6-3* were thoroughly analyzed and phenocopied *cap2*, both in terms of seed phenotypes and frequency. Moreover, *cap2, apc6-2* and *apc6-3* could be rescued by molecular complementation using an *APC6* transgene. In line with the gametophyte maternal effect phenotype, *APC6* was shown to be a maternally expressed imprinted gene. To this end, we could demonstrate that mutation of *APC1*, another subunit of the APC/C, displays a female gametophyte phenotype and enhances the frequency of disrupted endosperm development in an apc6 genetic background. Similar to *cap2*/*apc6, APC1* is imprinted and only expressed from the maternal allele, suggesting a maternal bias to the anaphase promoting complex (APC/C).

## Materials and methods

### Plant material and growth conditions

Wild-type (WT) accessions Columbia (Col-0), Landsberg *erecta* (L*er*-1), Tsushima (Tsu-1), C24 and T-DNA mutant lines were obtained from the Nottingham Arabidopsis Stock Centre (NASC; (Scholl et al., 2000)). Seedlings were grown on MS-2 (Murashige and Skoog medium with 2% sucrose) plates in a 16-h-light / 8-h-dark cycle at 22°C for 10 days prior to transferring to soil. Plants were further grown in a 16-h-light / 8-h-dark cycle at 18°C. T-DNA mutant plant lines *apc6-2* (SAIL_442_F11), *apc6-3* (SALK_008789), *apc1-2* (SALK_059826; g), SALK_002024C (*SRF5*), SALK_070429C (*SRF5*), SAIL_1280_D04 (*SRF5*), SALK_120562 (intergenic), SALK_033225 (intergenic), *cmt3-11* (SALK_148381; (Chan et al., 2006)), *met1-7* (SALK_076522; (Kanno et al., 2008)) and *drm1-2;drm2-2* (N16383; (Chan et al., 2006)) were in the Col-0 accession background (except *apc6-3*; Col-3). The triple *drm1-2*;*drm2-2*;*cmt3-11* mutant was crossed with the hemizygous *met1-7* mutant to generate a hemizygous *met1-7*;*drm1-2*;*drm2-2*;*cmt3-11* quadruple mutant. Methyltransferase mutants were maintained hemizygous (Mathieu et al., 2007). Endosperm marker lines used in crosses with *apc6-2, proAT5G09370>>H2A-GFP* (*EE-GFP*) and *proAT4G00220>>H2A-GFP* (*TE1-GFP*), were in Col-0 accession background (van Ekelenburg et al., 2023). For all crosses, closed flower buds were emasculated 2 days prior to crossing to avoid self-pollination.

### Microscopy

Crossed siliques were manually dissected and seeds were mounted on a microscopy slide in a clearing solution of glycerol and chloral hydrate as previously described (Grini et al., 2002). Microscopic analyses were performed using an Axioplan2 Imaging microscope equipped with a Zeiss Axio cam HDR camera. For fluorescent phenotypic analyses, seeds were mounted on a microscopy slide in two drops of tap water or 0.1 % Tween in 1x phosphate buffered saline (PBS; pH 7.4) or 30% glycerol with 20 µg/ml Propidium Iodide (PI). Seeds were analyzed for GFP fluorescence using an Andor DragonFly Spinning Disk confocal microscope with a 488 nm wavelength diode laser (Oxford) and imaged with a Zyla4.2 sCMOS camera (Oxford).

### DNA isolation and whole genome sequencing

Leaf and flower tissue from twelve individual *cap2/CAP2* (*cap2*mut) and twelve *CAP2/CAP2* (*cap2*wt) plants was collected in duplicate in a 2-ml round bottom tube containing a 5 mm metal bead and flash-frozen in liquid nitrogen. The frozen tissue was ground using a Retsch homogenizer for 1 min at 30 s^-1^. Genomic DNA (gDNA) was isolated as described in the ChargeSwitch gDNA Plant Kit (Invitrogen) manual and gDNA was eluted in 40 µl 10 mM Tris buffer (pH 8,5 at room temperature). Isolated gDNA was quality checked by Nanodrop and DNA concentrations were determined by Qubit dsDNA BR Assay Kit (Life Technologies). DNA libraries were prepared using the KAPA Hyper Kit (Roche) with 24 Unique Dual Indexes (Illumina) and sequenced 150 bp paired-end over two lanes on the Illumina HiSeq 4000 system.

### Identification of the *cap2* causative single nucleotide polymorphism

Sequencing reads were pre-processed using fastp with default settings and the overrepresentation analysis parameter to remove reads with low quality score, irregular GC content, short length, and sequencing adapters present (Chen et al., 2018). Trimmed reads were then quality checked with outputs from fastp to ensure trimming had been successful. Trimmed reads were mapped to the TAIR10 reference genome using bowtie2 (Langmead and Salzberg, 2012). Variant calling was performed on aligned and sorted BAM files using samtools mpileup and piped to bcftools to produce VCF files (Li et al., 2009). VCF files were converted to SHOREmap format and analyzed using SHOREmap backcross and then annotated to output tables and plots of backcross SNPs (Sun and Schneeberger, 2015).

### RNA isolation and cDNA synthesis

RNA was isolated from plant tissue (seeds or flowers) as described in the Spectrum Total Plant RNA Kit and On-Column DNaseI Digestion Set (Sigma Aldrich) manual. The tubes were shaken in a MagNA Lyser Instrument (Roche) at 7000 rounds per minute (rpm) for 15 sec, centrifuged at 13000 rpm in 4 °C for 15 sec and then placed at -20 °C for 2 min. This procedure was repeated three times before proceeding with the protocol. cDNA was synthesized from 2.5 µl RNA as described in the SuperScript III Reverse Transcriptase (Invitrogen) manual and cDNA was purified using Wizard SV Gel and PCR Clean-Up System (Promega).

### Single nucleotide polymorphism analysis

Single nucleotide polymorphisms (SNPs) between accessions were identified using GabiPD GreenCards (The GABI Primary Database) or Polymorph 1001 (1001 Genomes) and SNPs were verified by sequencing. Parental-specific expression based on these SNPs was determined by restriction digestion. For *APC1*, a SNP at position 6551 (G in L*er*-1; A in Col-0, Tsu-1 and C24) gains a restriction site for *AlwI* in the L*er*-1 accession. For *APC6*, a SNP at position 1297 (C in L*er*-1; G in Col-0, Tsu and C24) results in the loss of the restriction recognition site for *SacII* for the L*er*-1 accession. No restriction recognition site was identified covering the SNP at position 1516 (T in Col-0; A in C24), and therefore a recognition site for *BgIII* was introduced in the C24 accession using the Derived Cleaved Amplified Polymorphic Sequences (dCAPS) Finder 2.0 tool (Neff et al., 2002). The WT accessions Col-0, L*er*-1, Tsu-1 and C24 were crossed reciprocally, and L*er*-1 was crossed maternally to *met1-7/+, ddc/+* and *ddcm/+*. Siliques were dissected at 4 DAP using a stereomicroscope and seeds were harvested in MagNA Lyser Green Beads tubes (Roche) tubes were collected into liquid nitrogen. Seeds from three mother plants, four siliques each, were pooled for each biological replicate. RNA isolation and cDNA synthesis were performed as described above.

The SNP-containing DNA region was amplified by PCR with TaKaRa Ex Taq DNA Polymerase on 50 ng cDNA template using specific primers (STable 3) and PCR products were verified by sequencing. Restriction digestion was performed on 400 ng PCR product with 10 U of enzyme in a final volume of 25 µl. Equal amounts of digested fragments were analyzed using DNA-1000-LabOnChip and 2100 Bioanalyzer (Agilent Technologies).

### Molecular cloning and genotyping

Genotyping was performed using the KAPA3G Plant PCR kit (Roche) and gene- and T-DNA specific primers (STable 3). PCR products were sequenced by Eurofins Genomic and analyzed using Geneious software. To construct the *proAPC6:APC6-GFP* plasmid, a genomic fragment of the *APC6* gene, including 2 kb of the 5’-upstream sequence, was amplified by PCR and cloned between the AscI and SmaI sites of the pGFP_NOSG vector (Iwase et al., 2017). The resulting construct was then recombined into pGWB601 vector (Nakamura et al., 2010) using LR Clonase II. Genetic constructs were transformed into *Agrobacterium tumefaciens* C58 strain and introduced into *capulet2, apc6-3* and WT accession L*er*-1 using the floral dip method (Clough and Bent, 1998).

### RNA sequencing analysis

Raw reads from RNA Seq data were obtained from online resources (Bjerkan et al., 2020). Trimmed reads were obtained using Cutadapt version 1.18 (Martin, 2011), version TrimGalore 0.6.2 (Krueger, 2012) and mapped to the Araport11 CDS reference transcriptome (Cheng et al., 2017) using Bowtie version 2.3.5.1 (Langmead and Salzberg, 2012) with parameters *-- no-unal --no-mixed --no-discordant --sensitive --end-to-end -k 1*. Libraries were normalized using DESeq2 version 1.30.1 (Love et al., 2014). Statistical analysis was performed using ‘plyr’ version 1.8.6 (Wickham, 2011) and visualized using RStudio with ‘ggplot2’ version 3.3.5 (Wickham, 2016).

For the analysis of parental ratios, mapped reads were extracted using SAMtools version 1.9 (Li et al., 2009) and SNP counting was performed in Geneious Prime version 2021.1.1 (http://www.geneious.com/) using 0.2 as variant (L*er*-1) frequency threshold (except *AGL23* - variant frequency threshold). Reads from all timepoints were merged and aligned to the Col-0 reference sequence. SNP counting was performed to present an overview of reliable SNPs between L*er*-1 and Col-0 for each gene. The allele frequencies of each SNP for each timepoint (>10 coverage per SNP) were weighted for the coverage and visualized in Rstudio with ‘ggplot2’ version 3.3.5 (Wickham, 2016) and ‘ggpattern’ version 0.2.0 (Mike FC and Trevor L Davis, 2022).

### Image Analysis and Figure Preparation

Images were processed using Fiji (Schindelin et al., 2012). Figures were assembled in Adobe Illustrator 2022 (Adobe Systems Incorporated, San Jose, USA).

### Accession Numbers

All sequences generated in this study have been deposited to the National Center for Biotechnology Information Sequence Read Archive (https://www.ncbi.nlm.nih.gov/sra) with project number PRJNA808889.

### Supplemental data

All supplemental data files have been deposited to GitHub (https://github.com/PaulGrini/capulet2).

## Results

### *capulet2* causative mutation is located in *APC6*

The *capulet2* mutant was identified in an ethyl methanesulfonate (EMS)-induced gametophytic mutant screen by a segregation distortion assay using the multi marker chromosome 1 (mm1) line (Grini et al., 1999). The location of *cap2* was determined between the flanking genetic markers *ap1* and *gl2*

(Figure 1a) at a genome interval between *ADH1* and *gl2* using molecular markers (Grini et al., 2002). To determine the causative mutation of the *cap2* we employed whole genome sequencing. For this purpose, gDNA was isolated from twelve *cap2/CAP2* (*cap2/+*) and twelve *CAP2/CAP2* (+/+) plants. We performed a nucleotide polymorphism analysis between pairs of *cap2/*+ and *CAP2/CAP2* individuals, which revealed a clear linkage break on chromosome 1 (SFigure 1). We found between 100 and 180 candidate SNPs in the *ADH1* and *gl2* interval (Figure 1b, SData 1). Candidate SNPs were further selected by their base alignment quality score (BAQ ≥70) (Li, 2011), the presence of guanine to adenine conversion as the mutation arose in a population treated with EMS, an allele frequency between 0.25 and 0.75 (STable 1a) and conserved across all twelve comparisons (STable 1b). We found three SNPs that meet these requirements – one in the *ANAPHASE PROMOTING COMPLEX 6* (*APC6*), one in the *STRUBBELIG RECEPTOR FAMILY 5* (*SRF5*) and one in an intergenic region.

**Figure 1:**
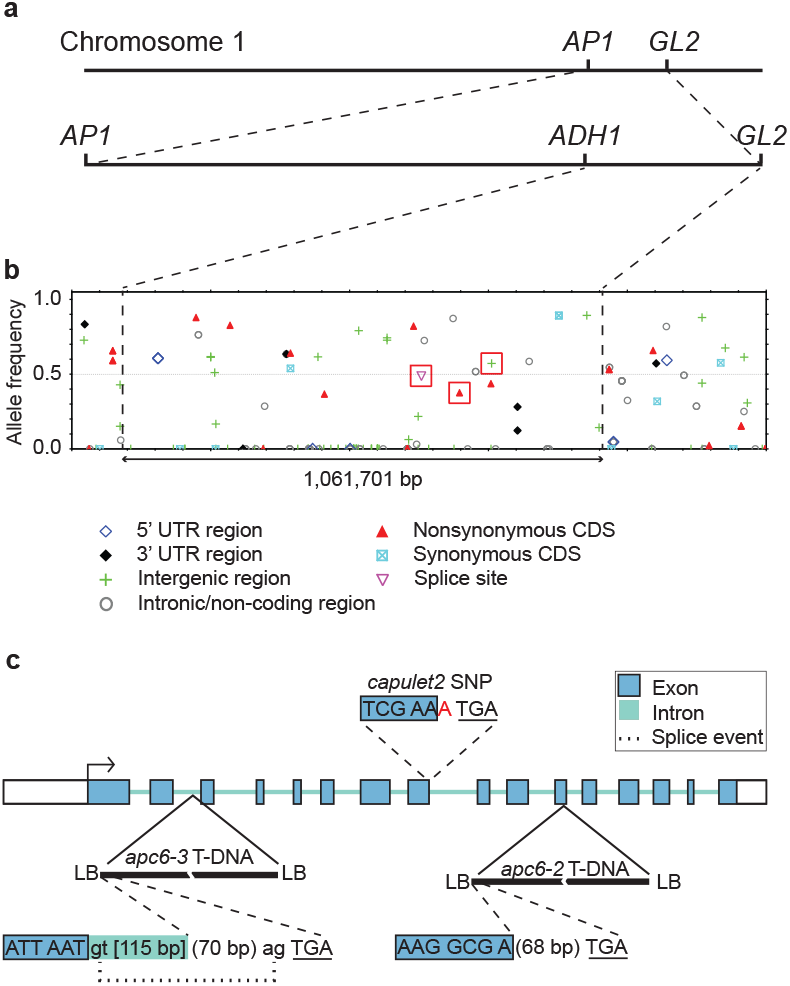
Identification of the causative *cap2* mutation. a) Identification of candidate causative SNPs located on chromosome 1 between the visible mutant markers *ap1* and *gl2* of multiply marked chromosome 1 (mm1). Segregation analysis using molecular markers narrowed down the location of the cap2 mutation between *ADH1* and *gl2*. b) SNP analysis of twelve independent comparisons between a *cap2*/*CAP2* and a *CAP2*/*CAP2* individual. Representation of identified SNPs, the type of mutation and allele frequency in a region of 1.06 Mbp of comparison 3. c) Molecular analysis of *cap2* and *apc* mutant alleles: The *cap2* G → A conversion (red) abolishes splicing and intron translation leads to a stop codon directly after the *cap2* SNP. The *apc6-2* T-DNA insertion, located in exon 12, has a 10 bp deletion and an in-frame stop codon 68 bp into the T-DNA. The *apc6-3* T-DNA insertion located in intron 2 has a 12 bp deletion. cDNA analysis revealed a cryptic splice acceptor site present in the T-DNA 70 bp inwards, resulting in splicing of intron 2 (up till the T-DNA insertion site) and part of the T-DNA, followed directly by an in-frame stop codon. Both *apc6-2* and *apc6-3* have at least two truncated T-DNA insertions at the same locus consecutively in opposite orientations. Exons and introns are highlighted by color and letter size (blue uppercase and green lowercase, respectively). Dotted line indicates splice event. Underlined nucleotides represent a stop codon. LB = T-DNA left border.

### *APC6* mutants are defective in endosperm development

We obtained three homozygous T-DNA insertion lines for *SRF5* and two T-DNA insertion lines flanking the intergenic candidate SNP and found that homozygous mutant plants for these insertions had normal seed sets and were indistinguishable from wild-types plants (*data not shown*). Our genome sequencing analysis revealed that the *cap2* mutant carries a splice site mutation in intron eight of *APC6*, potentially resulting in an unspliced transcript, introducing an in-frame early stop codon (Figure 1c). We obtained two T-DNA insertion lines for *APC6*, one that we named *apc6-2* (SAIL_442_F11) carrying an insertion located in exon twelve that results in the formation of a premature stop codon, and a second named *apc6-3* (SALK_008789) that carries an insertion located in the second intron, resulting in a cryptic splice acceptor site inside the T-DNA sequence leading to an in-frame stop codon (Figure 1c).

We found that when heterozygous *apc6-2* and *apc6-3* were crossed with wild-type (WT) pollen, the resulting seeds displayed developmental defects that were apparent at four days after pollination (DAP) and that resembled the seed phenotype observed in *cap2/+* (Figure 2a). To further characterize the endosperm phenotypes, we crossed *apc6-2/+* with an Early Endosperm (*EE-GFP*) reporter line expressing GFP before endosperm cellularization and a Total Endosperm (*TE1-GFP*) reporter line expressing GFP after endosperm cellularization (Bjerkan et al., 2023; van Ekelenburg et al., 2023). We found that a large proportion of seeds in *apc6-2/+*;*EE-GFP* plants had a reduced number and enlarged endosperm nuclei marked by a strong EE-GFP signal at 4 DAP (Figure 2a; SFigure 2). In *apc6-2/+*;*TE1-GFP* plants, however, the post-cellularization marker TE1-GFP was rarely detected in phenotypically mutant endosperm 4 DAP, suggesting that the mutant endosperm remain uncellularized (SFigure 2).

**Figure 2:**
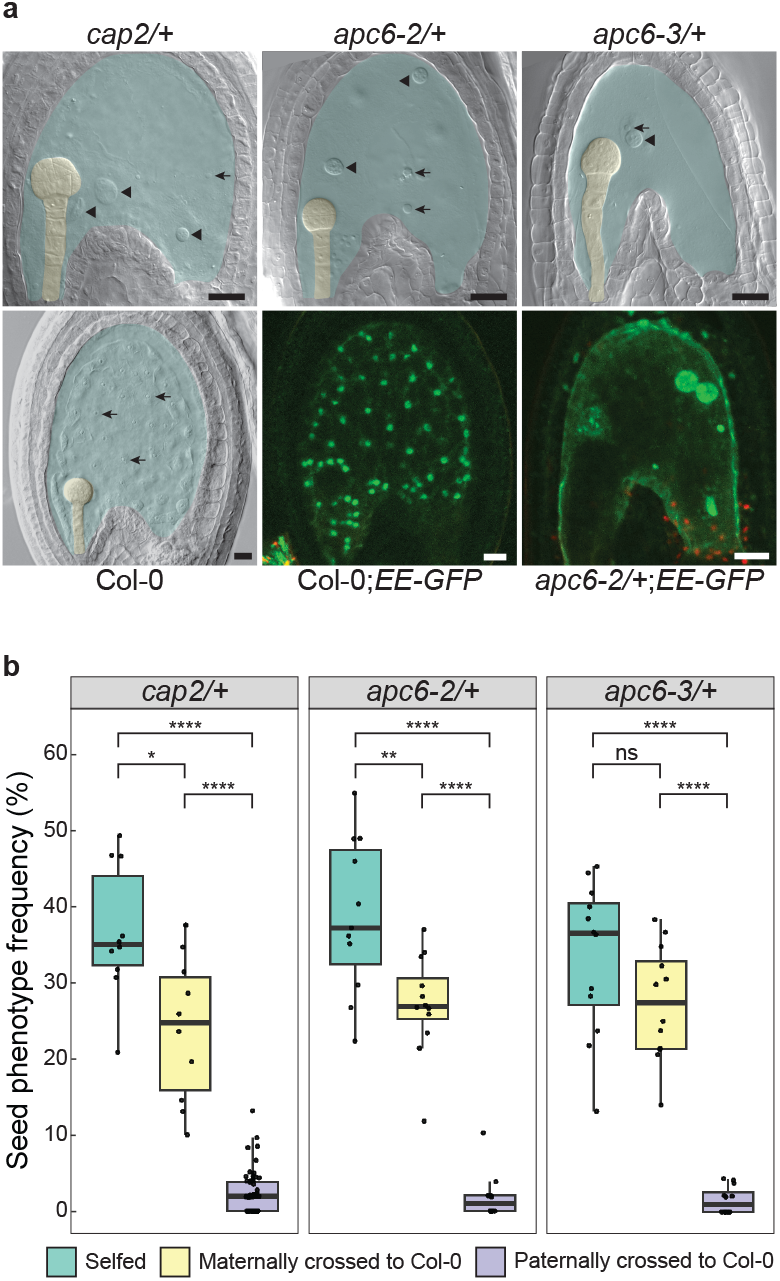
Seed phenotypes in *cap2, apc6-2* and *apc6-3* display gametophytic maternal effects. a) Endosperm phenotypes of *cap2/+, apc6-2/+* and *apc6-3/+* crossed maternally to Col-0 display arrested endosperm development at four days after pollination (DAP). A lower number of enlarged endosperm nuclei are observed compared to wild-type Col-0. Closed arrowheads indicate enlarged and abnormally shaped endosperm nuclei, arrows indicate smaller endosperm nuclei. A turquoise hue is added to highlight the endosperm, yellow hue highlights the embryo. Selfed *EE-GFP* compared to *apc6-2/+;EE-GFP* highlights enlarged and abnormally shaped endosperm nuclei. GFP-expressing seeds were counterstained with Propidium Iodide (red). Scale bar = 20 µm. b) Mutant seed phenotype frequencies for *cap2/+, apc6-2/+* and *apc6-3/+* at 4 DAP when selfed or crossed reciprocally to Col-0. n > 10 siliques per cross. Statistics was performed according to the Wilcoxon rank sum test; *** p-value < 0.001, **** p-value < 0.0001.

Notably, the *Arabidopsis nomega* mutant, caused by transposon insertion in *APC6* (Kwee and Sundaresan, 2003) and an *APC6* mutant in rice (Awasthi et al., 2012) have both been reported to be gametophytic mutants displaying pre-fertilization developmental defects. To investigate this discrepancy, we analyzed unfertilized ovules of wild-type, *cap2/+, apc6-2/+* and *apc6-3/+*. We found that the embryo sac central cell and egg cell, and egg apparatus of mutant lines are indistinguishable from WT (STable 2), and the three independent mutant alleles that we have identified for *APC6* show only post-fertilization seed defects.

We previously reported that *cap2* is a female gametophyte mutant, because seed defects are only observed when the mutant allele is transmitted maternally (Grini et al.,2002). We carried out reciprocal crosses between *cap2/+, apc6-2/+* and *apc6-3/+* and WT plants and quantified defects in seed development (Figure 2b, SData 2). We found that seeds failed to develop only when mutant lines were used as female parents (24.2%, 27.3% and 27.5% for *cap2/+, apc6-2/+* and *apc6-3/+* respectively), while seeds developed normally when wild-type plants were crossed with pollen from mutant lines. As initially reported for *cap2* (Grini et al., 2002), we did not find homozygous individuals in progenies of self-pollinated *apc6-2/+* and *apc6-3/+* plants (N = 47 and 42, respectively). Collectively, these findings indicate that mutations on *APC6* result in a maternal gametophyte effect on seed development.

### Genetic complementation of mutants with *APC6*

To verify that *cap2* is caused by mutation of *APC6*, we transformed *cap2/+* and *apc6-3/+* plants with a *proAPC6:APC6-GFP* (*APC6-GFP*) transgene. The transgene was also transformed into WT L*er*-1 plants to confirm endosperm expression (SFigure 3).

Plants from unique transformation events were genotyped for mutant alleles and *APC6-GFP*, and mutant seed phenotype frequencies were determined (SData 3). The frequency of mutant phenotypes was significantly reduced for the alleles tested, both in a hemizygous (*APC6-GFP/+*) or homozygous (*APC6-GFP/APC6-GFP*) transgene background (Figure 3; SFigure 4). Notably, homozygous *cap2* genotypes were identified in the T2 generation (Figure 3; SFigure 5a, b).

**Figure 3:**
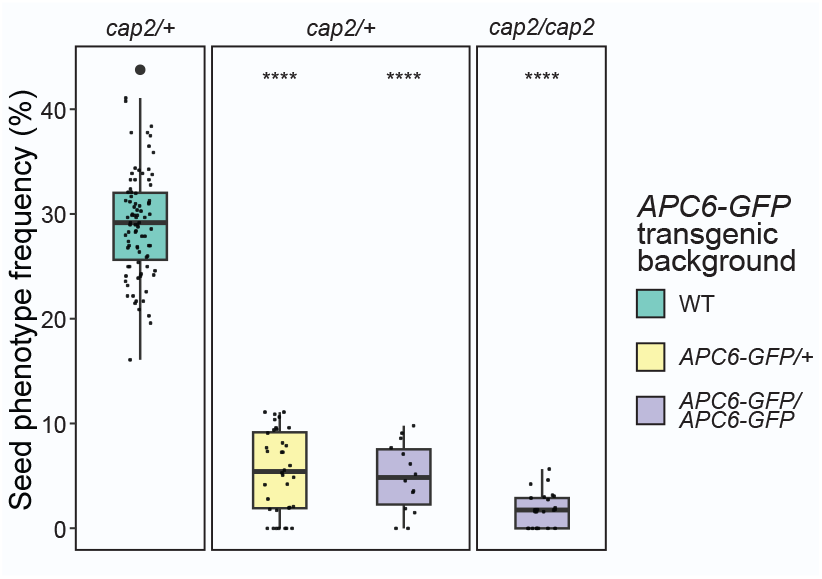
Complementation of *cap2* with an *APC6* transgene restores seed phenotype. Individuals heterozygous for the *cap2* SNP (*cap2/+*) were identified in both a hemizygous and homozygous transgenic background and homozygous individuals (*cap2*/*cap2*) were identified in a homozygous transgenic background. A significant reduction of the gametophyte maternal effect seed phenotype frequency is observed for *cap2/+* in a hemizygous transgenic background. In a homozygous transgenic background, a significant reduction is observed for *cap2/+* and *cap2*/*cap2*. Data from independent transgenic lines are merged for each box-plot: four independent rescue lines displayed significantly reduced phenotype frequency in a heterozygous *APC6-GFP* transgenic background, of which three segregated individuals with a homozygous construct background. Of these three lines, homozygous *cap2*/*cap2* mutants were identified in two. Statistics was performed according to the Wilcoxon rank sum test; **** p-value < 0.0001.

Several homozygous *apc6-3* individuals were detected in the T3 generation (SFigure 5c) and no mutant seed phenotypes were observed (SFigure 4). Homozygous *apc6-3* mutants in a homozygous rescue construct background crossed to *apc6-2*/+ plants, demonstrated several homozygous *apc6-2* individuals in the F2, as well as trans-homozygous *apc6-2*/*apc6-3* (SFigure 5d). Collectively, these data show that all *cap2/apc6* mutants can be rescued by a APC6-GFP transgene.

### Maternal penetrance and transmission of mutant alleles

We found that when plants carrying the three different *APC6* mutant alleles were pollinated with WT pollen, less than 50% of the seeds showed developmental abnormalities (Figure 2b), suggesting that these mutations were not fully penetrant. Therefore, we investigated in more detail the parental transmission of the different mutant alleles. To this aim, we selfed and crossed *cap2/+, apc6-2/+* and *apc6-3/+* reciprocally with WT plants and determined the transmission of the mutant alleles by molecular genotyping (*cap2* and *apc6-3*) or T-DNA herbicide resistance (*apc6-2*) of seedlings (Figure 4, SData 4). Paternal transmission of *cap2* (44.4%), *apc6-2* (47.4%) and *apc6-3* (46.7%) was not significantly different from the expected Mendelian transmission. Notably, the maternal transmission observed for *cap2* (30.5%), *apc6-2* (25.1%) and *apc6-3* (20.2%) was significantly lower than the expected 1:1 ratio when crossed to WT and also significantly lower than the expected 3:1 ratio in self crosses. These data support that maternal, but not paternal *cap2/apc6* alleles have significantly reduced transmission frequencies, suggesting a maternal bias in the role of *APC6*.

**Figure 4:**
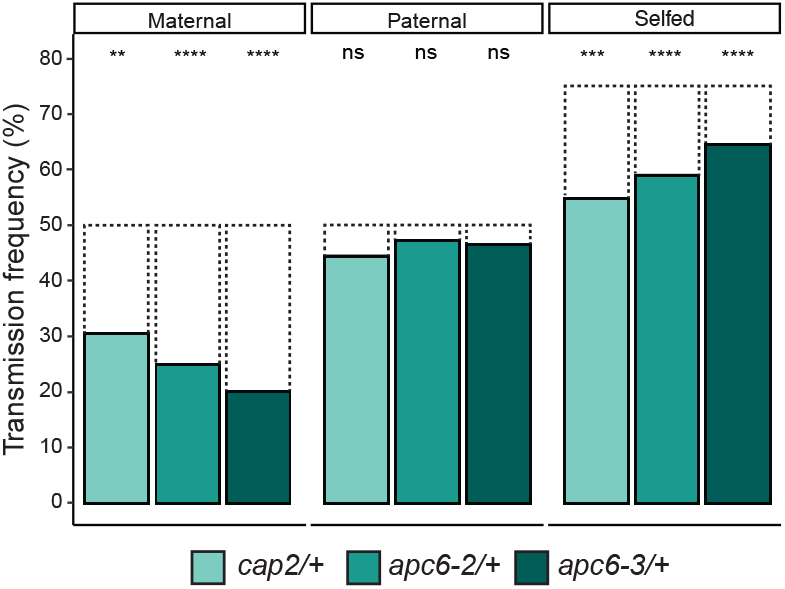
Genetic analysis shows reduced maternal transmission of *cap2, apc6-2* and *apc6-3*. Mutant alleles were selfed and crossed reciprocally to Col-0 and transmission was determined by genotyping (*cap2* and *apc6-3*) or using T-DNA herbicide resistance (*apc6-2*). Maternal transmission was significantly reduced while paternal transmission was not significantly different from the expected (50%, dashed line). Selfed transmission frequency for all mutant alleles was significantly lower than expected (75%, dashed line). Statistics was performed according to the Chi square test; ** p-value < 0.01, *** p-value < 0.001, **** p-value < 0.0001.

### Expression of *APC6* increases upon endosperm cellularization

We investigated *APC6* expression throughout seed development using publicly available RNA sequencing datasets (Bjerkan et al., 2020). As controls, we used several MADS-box transcription factors known to be expressed only in the endosperm, either bi-parentally or showing parental allelic bias (Colombo et al., 2008; Kang et al., 2008; Shirzadi et al., 2011). We found that *APC6* was highly expressed between 1 and 2 DAP, decreasing between three and 4 DAP and increasing again at 6 DAP (SFigure 6).

Since *APC6* shows a dynamic pattern of expression during seed development, we carried out a phenotypic characterization at different stages of seed development (2, 4, 6 and 9 DAP) using *cap2/+, apc6-2/+* and *apc6-3/+* plants crossed with wild-type Col-0 pollen. Generally, we found that both embryo and endosperm development was delayed compared to wild-type plants (SFigure 7) and in this phenotypic class endosperm proliferation was arrested after a few rounds of nuclear division. We also observed another phenotypic class where the endosperm was not fully cellularized at late stages in seed development (6 to 9 DAP), while embryo development was normal, although delayed when compared to wild-type plants (SFigure 7). The occurrence of seed phenotypes and collapsed seeds at 4 DAP and 6 DAP was highly similar in all *APC6* mutants (SFigure 7b, SData 5).

### *APC6* is a maternally expressed imprinted gene

Because all the *APC6* mutants identified showed maternal gametophyte effects on endosperm development, we hypothesized that this could be caused by uniparental expression of *APC6* transcripts. To test this hypothesis, we performed reciprocal crosses between different *Arabidopsis* accessions (L*er*-1, Col-0, C24 and Tsu-1) and analyzed the parental allele-specific expression of *APC6* in developing seeds by taking advantage of SNPs between these accessions (Figure 5a, SData 6). We found that at 4 DAP, the expression of *APC6* was exclusively from the maternal allele in most of the accessions tested. We quantified the maternal:paternal allelic ratios for the different reciprocal crosses and found that the maternal allele was consistently expressed between 8 and 16-fold higher than the paternal allele (Figure 5b, SData 6).

**Figure 5:**
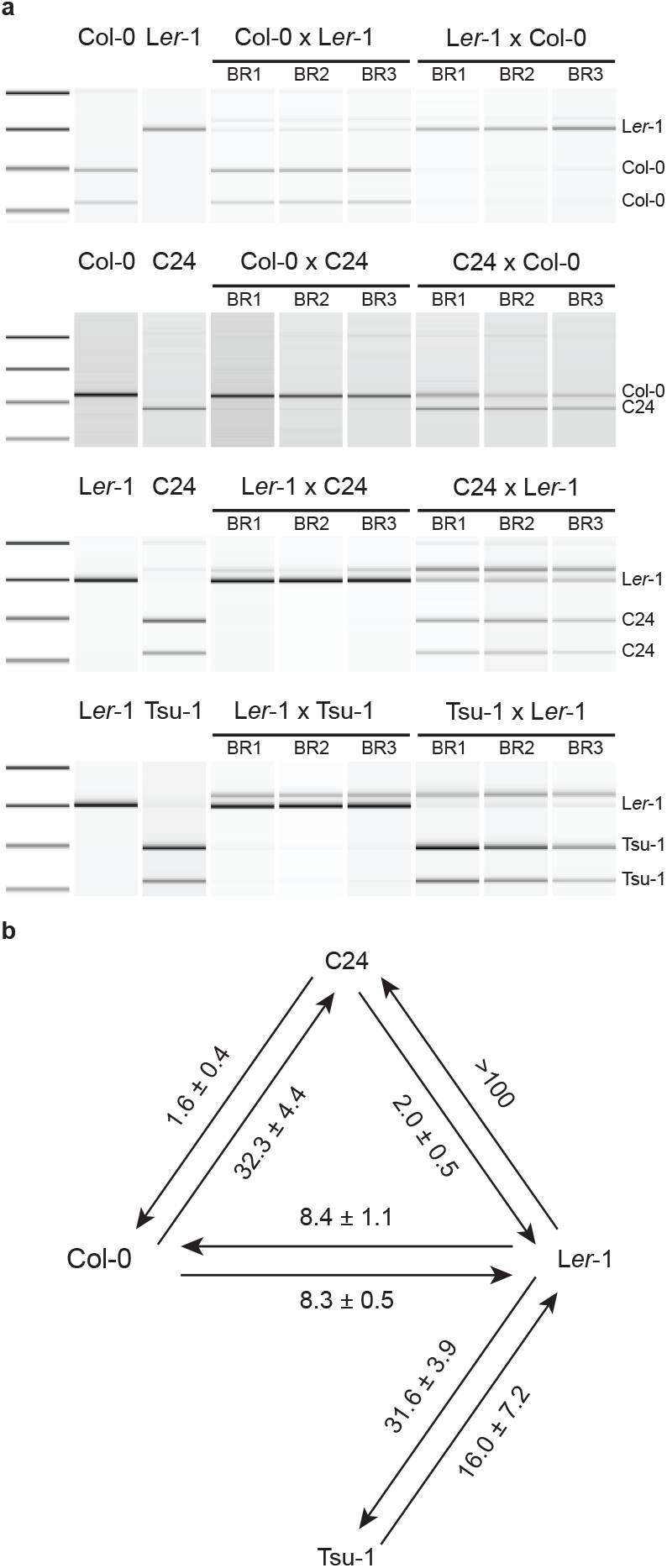
*APC6* is a maternally expressed imprinted gene. a) Imprinting analysis of *APC6* using accession-specific restriction digestion of SNPs in reciprocal crosses of Col-0, C24, L*er*-1 and Tsu-1. cDNA was generated from four days after pollination (DAP) crossed seeds. One replicate of homozygous wild-type controls is included as accession-specific cDNA reference and three biological replicates for each reciprocal cross are shown. One of two bioanalyzer replicates is presented and the accession of the maternal contributor is named first. In all directions, except for C24 as female parent, *APC6* is maternally expressed. Molarity ratios of maternal and paternal digested fragments from bioanalyzer profiles (a). Parental ratios are determined from all biological replicates and both bioanalyzer technical replicates. sd, standard deviation.

To verify these results, we analyzed RNA-seq data of early developing seeds (1 to 6 DAP) from reciprocal crosses between L*er*-1 and Col-0 (Shirzadi et al., 2011; Bjerkan et al., 2020). We identified several parental allele-specific SNPs, which we used to determine parental allele expression frequencies (Figure 6, SData 7). As controls we used MADS-box transcription factors, known to be expressed exclusively in the endosperm and biparentally (*AGL62*), maternally biased (*AGL36*) or paternally biased (*AGL23*) (Colombo et al., 2008; Kang et al., 2008; Shirzadi et al., 2011). We found that during early stages of seed development (1 to 4 DAP), *APC6* displays a strong maternal expression bias, similar to *AGL36*. The maternal bias of *APC6* was highly concordant with the molarity ratios determined experimentally (11.1-fold and 8.4-fold, respectively; SData7). However, whereas *AGL36* expression remains maternally biased at 6 DAP, the paternal expression of *APC6* increased, which could be attributed to an increasing contribution of the developing embryo.

**Figure 6:**
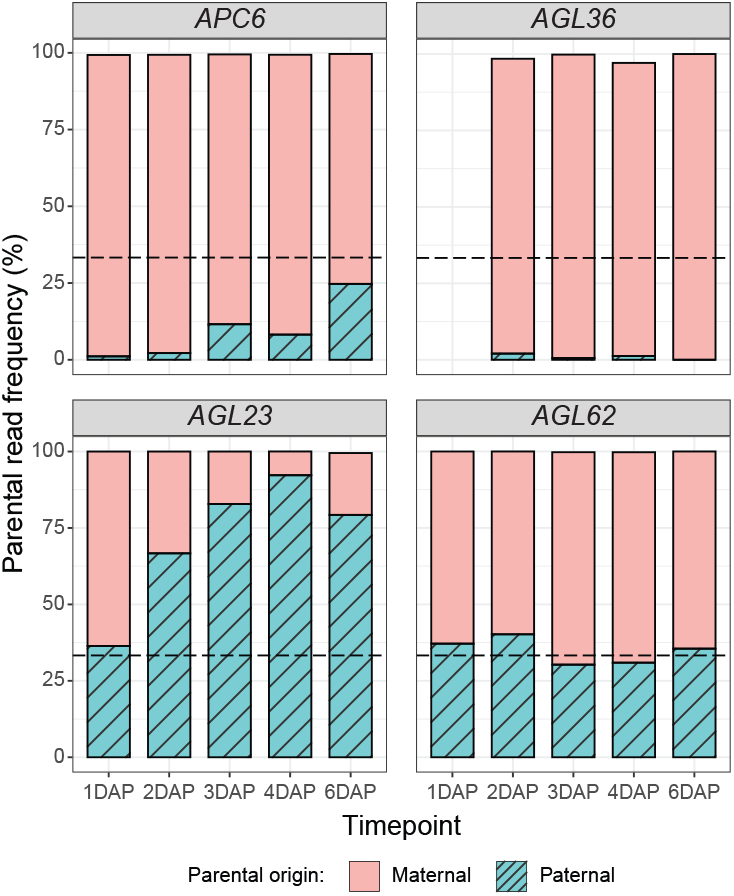
Seed stage analysis of parent specific allelic expression of *APC6* and imprinting control genes. Frequency of reads mapped to parental alleles of *APC6, AGL36* (MEG control), *AGL23* (PEG control) and *AGL62* (BEG control) at different timepoints (one, two, three, four and six days after pollination (DAP)). Maternal:paternal read frequency ratios are plotted at respective timepoints. SNP analysis was performed with Geneious Prime version 2021.1.1 using 0.2 as variant (L*er*-1) frequency threshold (except *AGL23* - 0.05 variant frequency threshold) and SNPs were weighted according to their coverage.

We hypothesized that the silencing of the paternal *APC6* allele may be caused by DNA methylation – a common characteristic of *Arabidopsis* MEGs (Gehring and Satyaki, 2017; Hornslien et al., 2019; Batista and Köhler, 2020). To test this hypothesis, we pollinated WT L*er*-1 plants with pollen from *met1-7/+* and *drm1-2*;*drm2-2*;*cmt3-11* (*ddc/+*) plants (SFigure 8a and b respectively). Due to possible DNA methyltransferase redundancy (Zhang and Jacobsen, 2006), we also generated a quadruple *met1-7*;*drm1-2*;*drm2-2*;*cmt3-11* (*ddcm/+*) mutant and used as pollen donor (SFigure 8c). Depletion of DNA methylation in the pollen, using single, triple or quadruple mutants did not reactivate the silenced paternal expression of *APC6* (SFigure 8a-c). Notably, the maternal:paternal expression ratios for all methyltransferase crosses were similar to the WT controls (Figure 5b, SFigure 8d).

Collectively, these data suggest that *APC6* is a maternally expressed imprinted gene and that imprinting of *APC6* is not mediated by canonical DNA methylation.

### Maternal expression of the APC/C

The APC/C is a large, multi-subunit complex consisting of at least 14 subunits in *Arabidopsis*. These subunits can be divided into groups based on their structure and function, such as scaffolding, catalytic, substrate recognition and platform units (Eloy et al., 2015). APC6 is a scaffolding unit consisting of tetratricopeptide repeats (TPRs), like APC3a and b, APC7 and APC8. In order to test whether members of other functional groups of the APC/C also display a maternal preference, we investigated *APC1*, a platform subunit containing proteasome-cyclosome (PC) repeats. APC1 is the largest subunit of the APC/C, and has previously been described to be important for female gametogenesis and embryogenesis (Wang et al., 2013).

Heterozygous *apc1-2/+* mutants displayed abnormalities in embryo sac formation, but no apparent reduction in seed set was found (SFigure 9a, SData 8). When *apc1-2/+* was crossed maternally to Col-0, gametophyte maternal effect endosperm phenotypes highly reminiscent to *cap2/apc6* were found, both in single *apc1-2/+* mutants and in an *apc1-2/+;apc6-2/+* double mutant (Figure 7a). Corresponding to mutants of *apc6, apc1-2/+* mutants displayed endosperm defects only when crossed maternally to Col-0, with a phenotype frequency of 37.1 % (Figure 7b).

**Figure 7:**
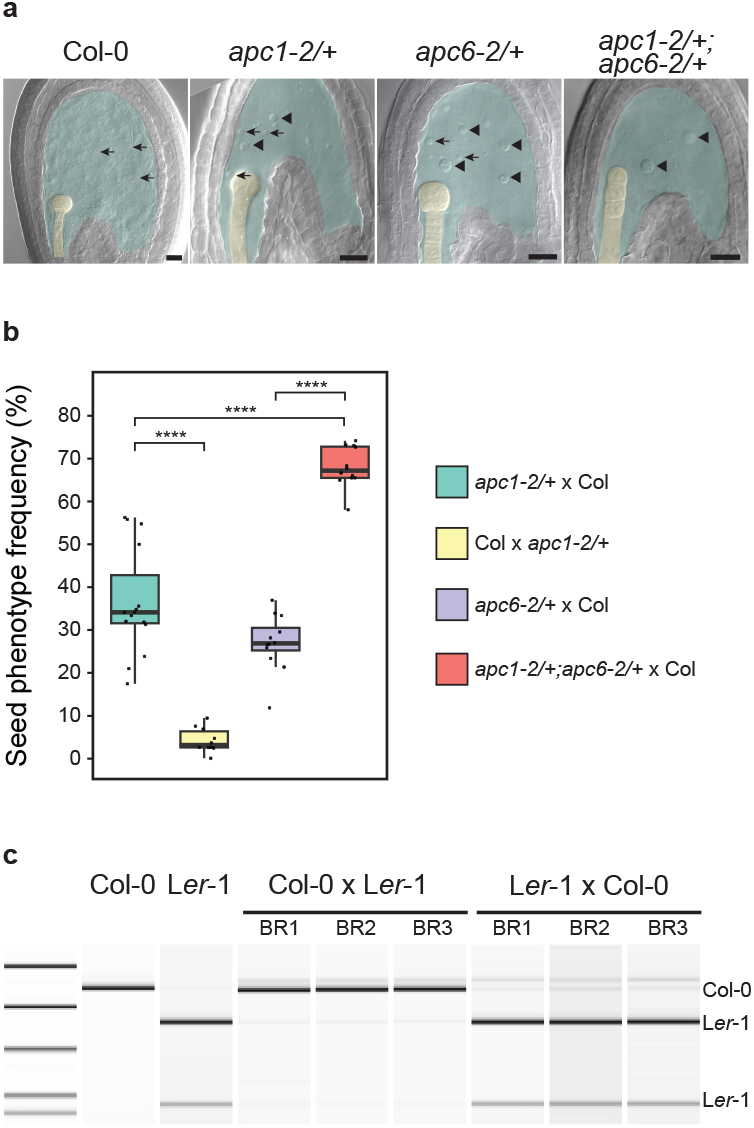
*APC1* mutants display a *cap2*/*apc6* endosperm phenotype and is a maternally expressed imprinted gene. a) Seed phenotypes of *apc1-2/+* and *apc1-2/+;apc6-2/+* double mutant at four days after pollination (DAP) when crossed maternally to Col-0. A turquoise hue is added to highlight the endosperm, yellow hue highlights the embryo. Arrows indicate small nuclei, arrowheads indicate large nuclei. Scale bar = 20 µm. b) Seed phenotype frequencies of *apc1-2/+* crossed reciprocally to Col-0 and *apc6-2/+* and *apc1-2/+;apc6-2+* crossed maternally to Col-0 at 4 DAP. No seed phenotype is present when *apc1-2/+* is crossed paternally, significantly different from when it is crossed maternally. The phenotype frequency of maternally crossed *apc1-2/+;apc6-2/+* double mutants is significantly increased compared to both single mutants. Statistics was performed according to Wilcoxon rank sum test; **** p-value < 0.0001. c) Imprinting analysis of *APC1* using accession-specific restriction digestion of SNPs in reciprocal crosses between Col-0 and L*er*-1 at 4 DAP. One replicate of each homozygous wild-type control and three biological replicates for the reciprocal crosses are shown. One of two bioanalyzer replicates is presented and the accession of the maternal cross partner is named first. Mostly maternal *APC1* expression was found in both directions of the reciprocal crosses shown.

When crossing the *apc1-2/+;apc6-2/+* double mutant maternally to Col-0 we observed a synergistic epistatic effect where the phenotype frequency of the double mutant (67.1 %) was significantly increased compared to both single mutants (Figure 7b). In addition, transmission of mutant alleles from a selfing *apc1-2/+;apc6-2/+* double mutant was investigated. Using the phenotype frequencies of *apc1-2/+* and the transmission frequency of *apc6-2/+*, we calculated the expected transmission frequencies to be 60.9 % for the *apc6-2* allele and 34.9 % for the *apc1-2* and *apc6-2* alleles together (SData 8). Notably, transmission of the *apc6-2* allele deviated from this expectancy, with a significant decrease in offspring carrying the mutant allele (57.4 %; SFigure 9b). However, the transmission of both mutant alleles together was not significantly different from our expectation.

Lastly, we investigated whether *APC1* expression was maternally biased by genomic imprinting. To this end, we analyzed the parental allele-specific expression of *APC1* in developing seeds using different *Arabidopsis* accessions (L*er*-1, Col-0, C24 and Tsu-1) as described previously. In 4 DAP seeds, *APC1* was exclusively expressed from the maternal allele in all accessions and reciprocal crosses (Figure 7c, SFigure 9c).

Collectively, these data suggest that both *APC1* and *APC6* are maternally expressed imprinted genes that are required maternally both in the scaffolding and platform units of the APC/C for early endosperm development, suggesting a maternal bias in the role of the APC/C complex.

## Discussion

In this study, we identified that mutation in *APC6* is associated with the female gametophyte maternal effect mutant *capulet2* (*cap2*). Molecular characterization of *cap2* has revealed that *APC6* is a maternally expressed imprinted gene implicated in endosperm proliferation and cellularization. Our data reveals that the maternal contribution of the APC/C is critical for endosperm development in plants.

### *capulet2* causative mutation is located in *APC6*

We have found that *cap2* is caused by a splice donor mutation in *APC6*. We were able to functionally complement *cap2, apc6-2* and *apc6-3* mutants with an *APC6* transgene, allowing the generation of homozygous *cap2, apc6-3* and *apc6-2/apc6-3* mutant plants. Previous studies have revealed that mutations on several APC/C subunits result in female gametophyte defects (Saleme et al., 2021), including *Arabidopsis nomega* that carry a transposon insertion in *APC6* (Kwee and Sundaresan, 2003) and a mutant in rice *OsAPC6* (Awasthi et al., 2012). Notably, we have found that *cap2, apc6-2* and *apc6-3* mutations only display defects on early seed development. We found that the female gametophyte of *cap2, apc6-2* and *apc6-3* was indistinguishable from wild-type.

### *APC6* is a maternally expressed imprinted gene

Our analysis, using both allele-specific assays and RNA-seq allele-specific expression quantification has revealed that *APC6* is an imprinted, maternally expressed gene. However, since we used whole seeds in the analysis, transcripts from the maternal seed coat or embryo could affect this interpretation (Schon and Nodine, 2017). To address this caveat, we analyzed publicly available expression data from early developing seeds (Belmonte et al., 2013), which showed that APC6 is primarily expressed in the endosperm. Due to the different genotypes of endosperm, embryo and seed coat (2:1; 1:1; 2:0, maternal:paternal respectively), a whole seed maternal:paternal ratio of ≈ 2:1 can be estimated if APC6 is equally expressed from both parental alleles. The maternal:paternal ratios in our analyses are at least >4-fold higher, ranging up to >50-fold higher, indicating that expression of APC6 is maternally biased in the endosperm. Our data also indicates that imprinting of APC6 is accession-specific, a phenomenon that also has been observed in previous studies of imprinted genes (Wolff et al., 2011; Pignatta et al., 2014).

It has been proposed that MEGs are regulated by DNA methylation through the activity of MET1 that silences gene expression in gametes and by DEMETER (DME) that removes methylation in the central cell and endosperm restoring the expression of maternal alleles (Choi et al., 2002; Xiao et al., 2003; Gehring et al., 2006). Here, we have investigated the role of DNA methylation in imprinting of *APC6* by crossing L*er*-1 with various DNA methyltransferase mutants in a Col-0 background and performed SNP digestion analysis to determine if expression of the paternal allele is reactivated. The single, triple and quadruple mutants of *met1-7, drm1-2*;*drm2-2*;*cmt3-11* and *met1-7;drm1-2*;*drm2-2*;*cmt3-11* were used to reduce redundancy by other methyltransferases. None of the mutant combinations showed an increased paternal expression and thus we concluded that imprinting of *APC6* is not caused by these canonical DNA methyltransferases.

This is consistent with previous reports where the majority of MEGs was not regulated by MET1 DNA methylation maintenance (Hornslien et al., 2019). Further analysis of the epigenomic landscape of *APC6* is thus required to determine if *APC6* imprinting is guided by histone guided DNA methylation or other histone modifications.

It is experimentally demanding to distinguish *de novo* transcripts after fertilization from transcripts originating from the central cell as maternal carry-over (Evans and Kermicle, 2001). The *APC6* expression profile is constructed from whole seed RNA (SFigure 5), and RNA originating from the seed coat or the embryo is present as discussed above. Compared to the endosperm-specifically expressed *AGL36, AGL23* and *AGL62*, overall expression of *APC6* is higher at the early stages (1 and 2 DAP) of seed development and carry-over of transcripts from the maternal central cell, as described on a general basis (Luo et al., 2014) cannot be excluded. However, at 4 DAP, allele-specific expression is mostly maternal, supporting the finding that *APC6* is imprinted. Long-lived mRNAs in the endosperm have been described for endosperm maturation phase transcripts (Matilla, 2022), but it seems unlikely that transcripts, originating from the maternal central cell, are abundant at the globular embryo stage. Moreover, microarray data indicated that expression of *APC6* at the globular embryo stage is higher in the endosperm compared to the seed coat and embryo (Belmonte et al., 2013). Additionally, the increase in paternal expression at 6 DAP could be due to a stronger impact of the growing embryo, as this coincides with consumption of the endosperm (Nowack et al., 2010; Lafon-Placette and Köhler, 2014). As such, late-stage expression of *APC6* is directed towards a parental expression ratio closer to the 1:1 diploid genotype of the embryo. Altogether, the findings of this study indicate that *APC6* is a maternally expressed imprinted gene in the endosperm.

### The APC/C is maternally biased

The APC/C is a large protein complex consisting of at least fourteen core subunits and essential for gamete development (Saleme et al., 2021). Genetic analyses have revealed that several subunits of the APC/C are essential for megagametogenesis (*APC2, APC3a/APC3b* and *APC10*; (Capron et al., 2003; Pérez-Pérez et al., 2008; Eloy et al., 2011)) and male gametogenesis (*APC8* and *APC13*; (Saze and Kakutani, 2007; Zheng et al., 2011)). Furthermore, overexpression of *APC8* displayed reduced seed set, suggesting that careful dosage regulation of *APC* subunits is essential for seed development (Zheng et al., 2011). Here we have demonstrated that both *APC6* and *APC1* are imprinted and expressed predominantly from the maternal genome. Both APC/C loci lead to a gametophyte maternal effect phenotype when mutated, and double mutants suggest an additive effect on seed survival. The strong maternal expression bias of *APC1* in all tested ecotypes further emphasize the importance of maternal control of the APC/C during early seed development. Interestingly, and in addition to *APC6* and *APC1* mutants, *APC4* and *APC11* also show gametophyte maternal effect phenotypes (Kwee and Sundaresan, 2003; Wang et al., 2012; Wang et al., 2013; Guo et al., 2016), in line with the notion of APC/C being under maternal control.

It remains unclear why subunits of such a crucial protein complex are subjected to parental allele-specific regulation, and the function of genomic imprinting in this regard remains to be determined. However, our data supports the gene dosage hypothesis for the evolution of genomic imprinting, which states that imprinting evolved as a mechanism to control and regulate gene expression levels (Dilkes and Comai, 2004; Ferguson-Smith, 2011). For subunits of the APC/C, where too little or too much expression of a gene is detrimental for seed development, genomic imprinting could be a measure to balance gene expression contributing to this protein complex. Taken together, our observations propose a maternal bias to the control of endosperm proliferation.

## Supporting information

STable 1

Stable 2

Stable 3

SFigure 1

SFigure 2

SFigure 3

SFigure 4

SFigure 5

SFigure 6

SFigure 7

SFigure 8

SFigure 9

## Author contributions

P.E.G. designed the research; Y.S.vE., I.V.M., M.P.H., S.K. & C.M. performed the experiments; Y.S.vE., I.V.M., C.M., J.G-M & P.E.G. analyzed and discussed the data; Y.S.vE., I.V.M., J.G-M & P.E.G. wrote the article; All authors revised and approved the article.

## Supporting information

The following supplemental materials are available.

**SFigure 1:** Distribution of SNPs between *cap2*/*CAP2* and *CAP2*/*CAP2*.

**SFigure 2:** GFP expression in *apc6-2/+* with endosperm marker lines.

**SFigure 3:** Verification of the *APC6* transgene functionality using confocal microscopy.

**SFigure 4:** An *APC6* transgene can complement the *apc6-3/+* mutant phenotype.

**SFigure 5:** Identification of homozygous *cap2, apc6-3* and *apc6-2* plants after complementation with an *APC6* transgene.

**SFigure 6:** Relative expression of *APC6* throughout seed development.

**SFigure 7:** Seed development in *APC6* mutants and wild-type.

**SFigure 8:** Imprinting of *APC6* is not regulated by canonical DNA methylation.

**SFigure 9:** Characterization of *apc1-2* ovules, *apc1-2/+;apc6-2* mutant allele transmission and maternal expression of *APC1*.

**STable 1:** Identification of the candidate SNPs for *capulet2*.

**STable 2:** Phenotypic characterization of the embryo sac.

**STable 3:** Primer name, sequence and comments.

**SData 1:** SNP characterization comparison

**SData 2:** Phenotype frequencies of *capulet2, apc6-2* and *apc6-3*

**SData 3:** Phenotype frequencies of rescued *capulet2* and *apc6-3*

**SData 4:** Transmission frequency *capulet2, apc6-2* and *apc6-3*

**SData 5:** Extended phenotype frequencies of *capulet2, apc6-2* and *apc6-3*

**SData 6:** Bioanalyzer data for *APC6* in WT and methyltransferase mutants

**SData 7:** Parental allele frequencies in *APC6* and control genes

**SData 8:** Genetic and phenotypic characterization of *APC1*

