## Supplementary material for "Molecular characterization of *capulet2* reveals the importance of *ANAPHASE PROMOTING COMPLEX 6* maternal expression in endosperm development": STable 1

**a**

| Comparison[1] | Total SNPs | ≥70 BAQ[2] | G > A[3] | Allele frequency[4] |
|---|---|---|---|---|
| 1 | 167 | 24 | 10 | 7 |
| 2 | 129 | 18 | 9 | 8 |
| 3 | 150 | 24 | 11 | 9 |
| 4 | 173 | 13 | 9 | 7 |
| 5 | 145 | 11 | 9 | 4 |
| 6 | 152 | 13 | 8 | 5 |
| 7 | 110 | 16 | 12 | 8 |
| 8 | 144 | 16 | 11 | 7 |
| 9 | 133 | 12 | 5 | 4 |
| 10 | 128 | 18 | 9 | 5 |
| 11 | 112 | 11 | 8 | 8 |
| 12 | 108 | 14 | 8 | 8 |

**b**

| Location | Type of mutation | AGI | Gene |
|---|---|---|---|
| 29,619,446 | Splice donor site change | AT1G78770 | *ANAPHASE PROMOTER COMPLEX SUBUNIT 6* (*APC6*) |
| 29,708,804 | CDS; nonsynonymous | AT1G78980 | *STRUBBELIG-RECEPTOR FAMILY 5* (*SRF5*) |
| 29,783,917 | Intergenic | - | - |
