## Supplementary material for "Molecular characterization of *capulet2* reveals the importance of *ANAPHASE PROMOTING COMPLEX 6* maternal expression in endosperm development": Stable 2

| Plant | Mature embryo sac | NA | Non-developed | Total |
|---|---|---|---|---|
| Col-0 | 81% | 17% | 2% | 185 |
| *cap2/+* | 81% | 17% | 2% | 161 |
| *apc6-2/+* | 84% | 16% | - | 97 |
| *apc6-3/+* | 77% | 20% | 3% | 187 |
