## Supplementary material for "Molecular characterization of *capulet2* reveals the importance of *ANAPHASE PROMOTING COMPLEX 6* maternal expression in endosperm development": Stable 3

| Primer | Oligo sequence (5' –> 3') | Comments |
|---|---|---|
| LBa1 | TGGTTCACGTAGTGGGCCATCG | Genotyping Salk T-DNA left border for *apc6-3*/*drm1;drm2/apc1-2* |
| LBb1.3 | ATTTTGCCGATTTCGGAAC | Genotyping Salk T-DNA left border for *met1-7*/*cmt3-11* |
| LB3 | TAGCATCTGAATTTCATAACCAATCTCGATACAC | Genotyping Sail T-DNA left border for *apc6-2* |
| APC6_SAIL442_F11_Fp | TGCCAACTCTGTACATTGGAATGGAG | Genotyping genomic forward for *apc6-2*; Sail T-DNA left/right border for *apc6-2* |
| APC6_SAIL442_F11_Rp | GACCACAGTTGGTTCCCAGG | Genotyping genomic reverse for *apc6-2* |
| APC6_SLK008789Nw_Fp | TGATGCTAAAGACGGGAACGTGA | Genotyping genomic forward for *apc6-3*; Salk T-DNA left border for *apc6-3* |
| APC6_SALK_008789_Rp | AGCACAGAGGATCAGCCTTGA | Genotyping genomic reverse for *apc6-3* |
| APC6_intron1-3_Rp | CACAGAGGATCAGCCTTGATAGCA | Genotyping Salk T-DNA right border *apc6-3* |
| APC6_intron7-9_Fp | GTGGTAGTGTCTCTGGCTCATCT | Genotyping forward for *capulet2* |
| APC6_3DOWN_2_Rp | GTAACAGTCTCTTGGTTATTGATGCATGGA | Genotyping reverse for *capulet2* |
| DRM2_SP | AGTTAGCCCGAGGGCCACCTTT | Genotyping genomic *drm1;drm2* |
| DRM2_ASP | CGCCTGCCAGATTGTTACAAGG | Genotyping genomic *drm1;drm2*; Salk T-DNA left border for *drm1;drm2* |
| cmt3_SALK_148381_ASP | CGTGAAGGTTGGCATATTCTTCTG | Genotyping genomic *cmt3-11*; Salk T-DNA left border for *cmt3-11* |
| cmt3_SALK_148381_SP | AGCTATGTCGACAGGGTTGTGC | Genotyping genomic *cmt3-11* |
| met1_SALK076522_Rp | CAGAACAGGTTTCCACCAAGG | Genotyping genomic *met1-7*; Salk T-DNA left border for *met1-7* |
| met1_SALK076522_Lp | TCAATGAGCAGTGGAAGCAAG | Genotyping genomic *met1-7* |
| APC6_ecot_SNP1_Fp | GCTGAGTACTACCATCAATG | *APC6* L*er*-1 x Col-0/Tsu-1/C24 SNP digestion forward primer |
| APC6_ecot_SNP1_Rp | CACGCCATTAGATAGAGC | *APC6* L*er*-1 x Col-0/Tsu-1/C24 SNP digestion reverse primer |
| ColC24_Fp | CTTGGTTTGCGGTGGGTTG | *APC6* Col-0 x C24 SNP digestion forward primer |
| ColC24_BglII_Rp | CGAGCTGGTGAGAATGAGAGATC | *APC6* Col-0 x C24 SNP digestion reverse primer |
| APC6-F | TGGCGCGCCCGAGGTATGGATAAAACTCTATCTCCGT | Amplifying *APC6* with promoter for construction of *proAPC6:APC6:GFP* |
| APC6-R | AATCCCGGGGCAGAGCTCAACCTTTGAATCAACCCCG | Amplifying *APC6* with promoter for construction of *proAPC6:APC6:GFP* |
| apc1-2_Fp | CTTGGTCGAGGGACAGAAGC | Genotyping genomic forward for *apc1-2* |
| apc1-2_Rp | CAGGTTCCTCTAGTGGATCTCTTAGAATCAG | Genotyping genomic reverse for *apc1-2;* Salk T-DNA |
