## Supplementary material for "Molecular characterization of *capulet2* reveals the importance of *ANAPHASE PROMOTING COMPLEX 6* maternal expression in endosperm development": SFigure 1

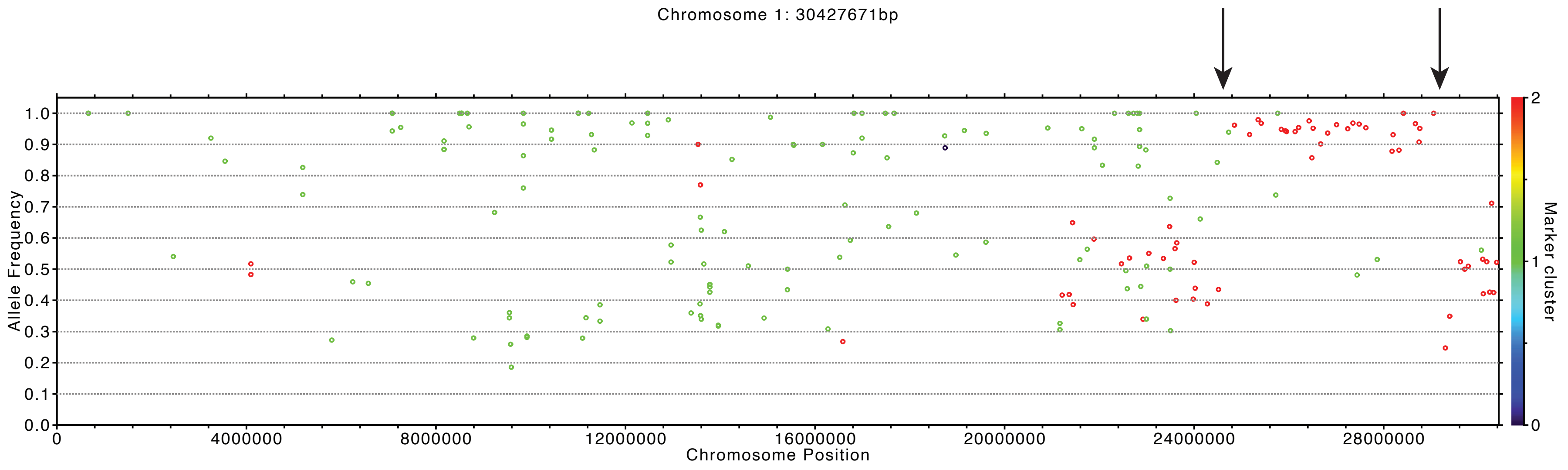


*Points in colors: AF (=alt divided by (alt+ref)) at markers with clustering. alt or ref: coverage of non-reference or reference allele
*rank/cluster table: min_fg_cov max_fg_cov fg_qua bg_mut_cov bg_mut_qua size_cluster

cluster 2: (L:36.9, H:77.1) 220.2 0.0 0.0 63
cluster 1: (L:29.0, H:85.0) 67.9 0.0 0.0 132
cluster 0: (L:48.0, H:66.0) 42.0 40.0 88.0 1
