## Supplementary figures and images for "Molecular characterization of *capulet2* reveals the importance of *ANAPHASE PROMOTING COMPLEX 6* maternal expression in endosperm development"

### SFigure 2

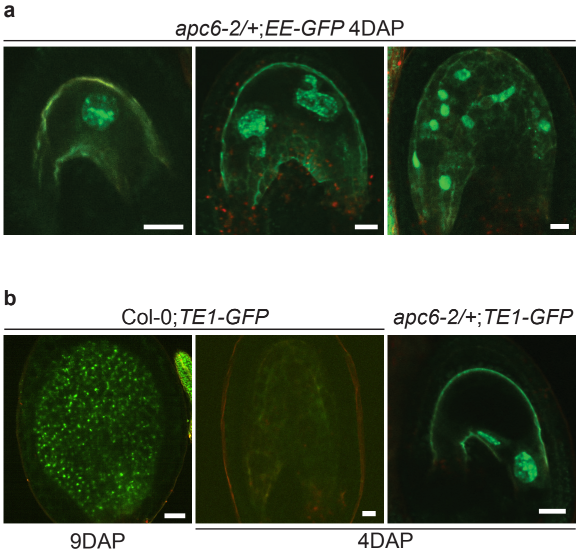

### SFigure 3

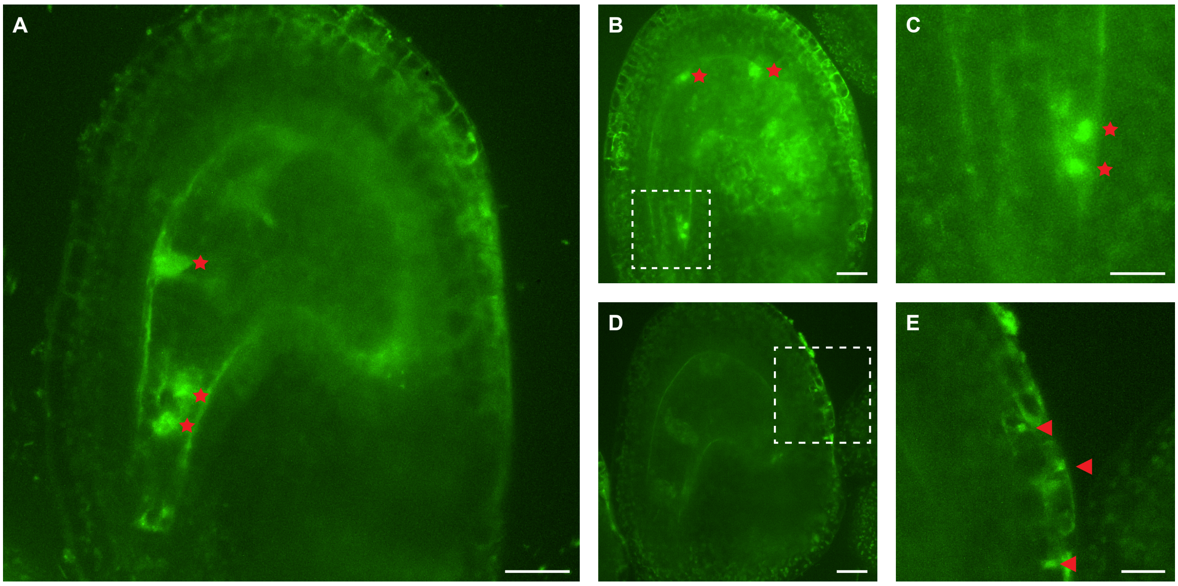

### SFigure 4

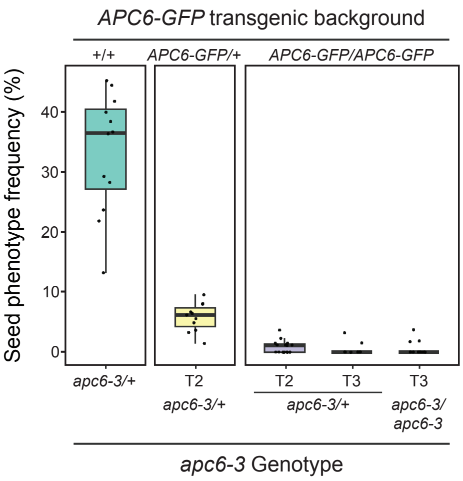

### SFigure 5

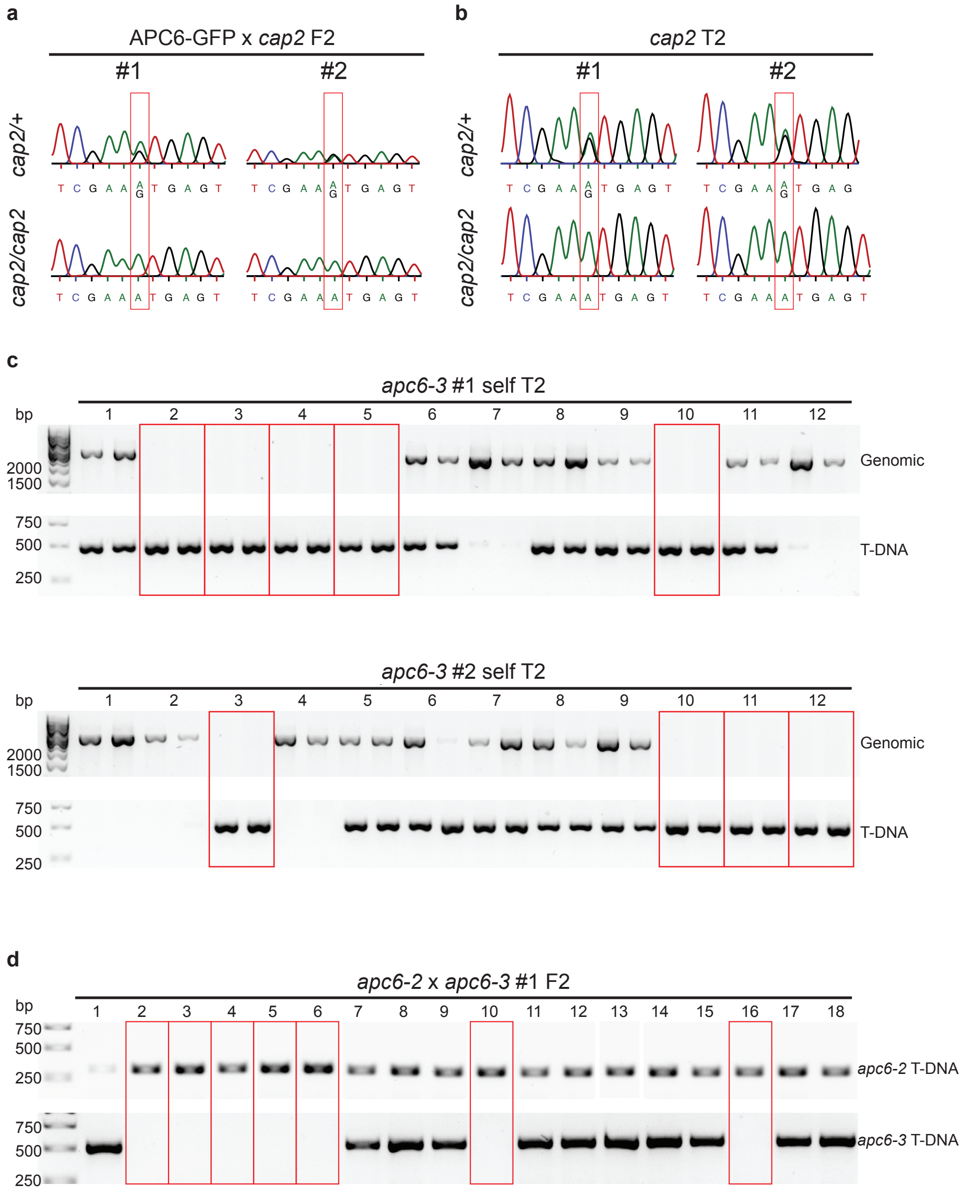

### SFigure 6

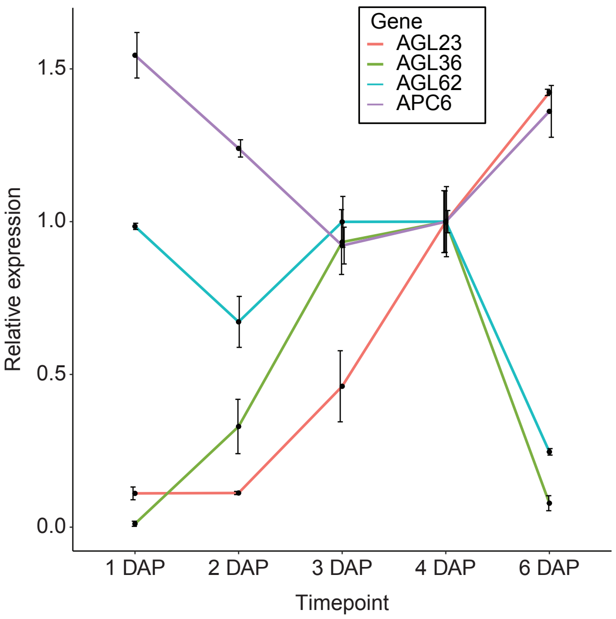

### SFigure 7

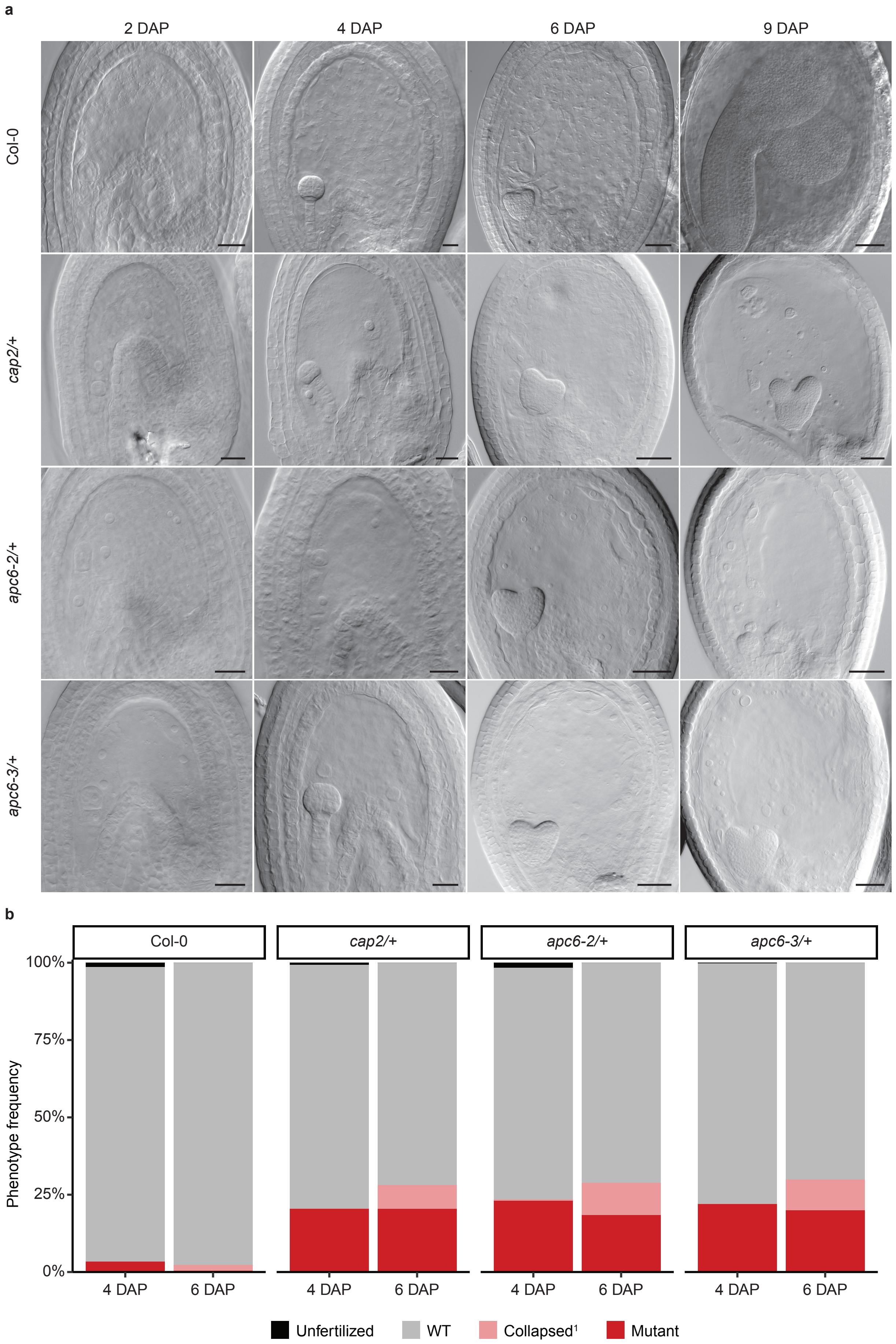

### SFigure 8

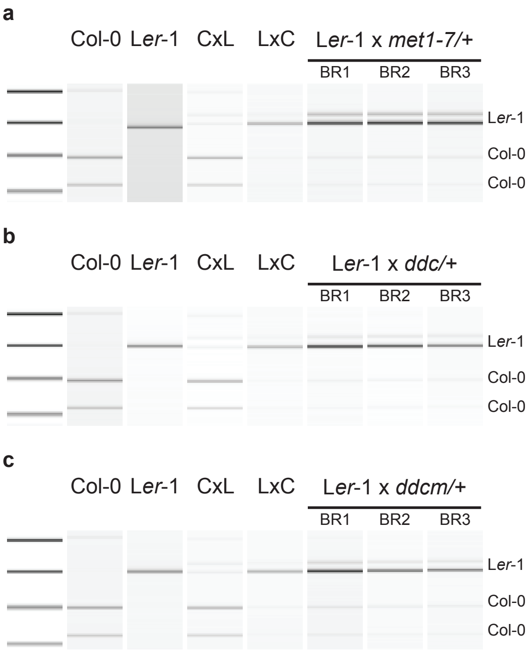


**d**

| Cross | m:p ratio |
|---|---|
| L*er*-1 x *met1-7/+* | 9.1 ± 1.0 |
| L*er*-1 x *ddc/+* | 9.8 ± 2.1 |
| L*er*-1 x *ddcm/+* | 7.9 ± 0.5 |

### SFigure 9

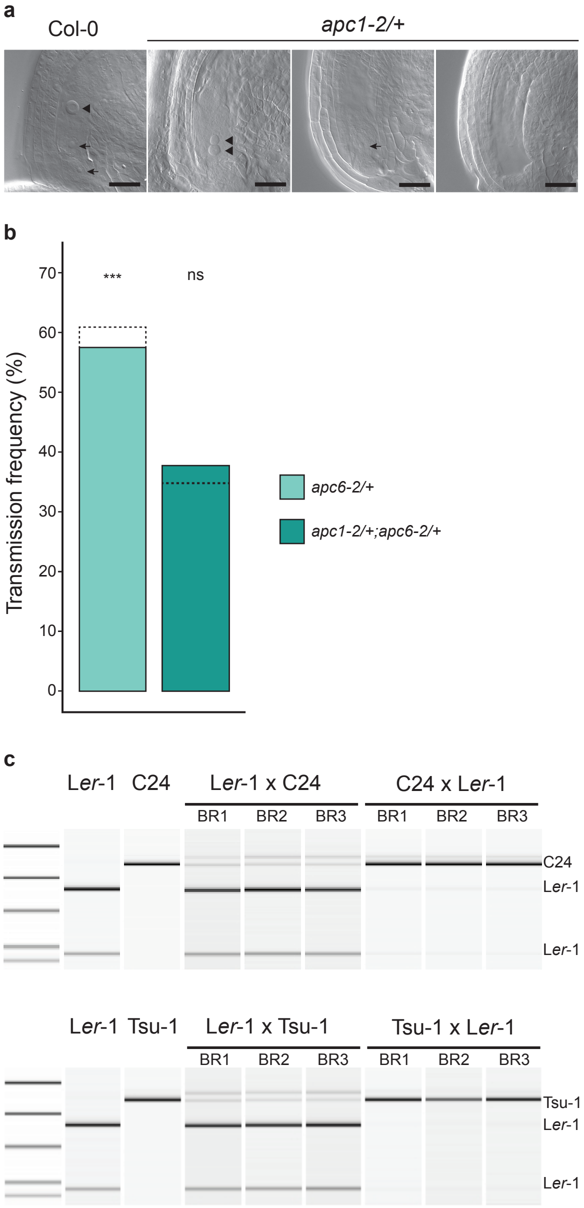
